# Differences in Chemo-signaling Compound-Evoked Brain Activity in Male and Female Young Adults: A Pilot Study in the Role of Sexual Dimorphism in Olfactory Chemo-Signaling

**DOI:** 10.1101/2020.09.03.280685

**Authors:** Taylor D. Ottesen, Kevin C. Davis, Landon K. Hobbs, Nathan M. Muncy, Nicholas M. Stevens, Malia Anderson, Paula Johnson, Christopher R. Doxey, Kent Richter, Haonan Wang, Randy Hartley, C. Brock Kirwan, Jonathan J. Wisco

**Affiliations:** Brigham Young University, Department of Physiology and Developmental Biology, Neuroscience Center, Provo, UT 84602, USA; Yale School of Medicine, New Haven, CT 06519, USA; University of Miami Miller School of Medicine, Miami, FL 33136; University of Virginia School of Medicine, Charlottesville, VA 22908, USA; Brigham Young University, Department of Exercise Science, Neuroscience Center, Provo, UT 84602, USA; Touro University Nevada College of Osteopathic Medicine, Henderson, NV 89014, USA; Utah Valley University, Orem, UT 84058, USA; Department of Mechanical Engineering, Neuroscience Center, Brigham Young University, Provo UT; Department of Psychology, Neuroscience Center, Brigham Young University, Provo UT; Mayo Clinic School of Medicine, Scottsdale, AZ 85259, USA; Department of Electrical and Computer Engineering, Brigham Young University, Provo UT; University of Utah School of Medicine, Department of Neurobiology and Anatomy, Salt Lake City, UT 84132, USA; Boston University School of Medicine, Department of Anatomy and Neurobiology, Boston, MA 02118

**Keywords:** Sexual Dimorphism, Olfactory Chemo-Signaling, Pheromones, 4,16-androstadien-3b-ol, 1,3,5(10),16-Estratetraen-3-ol, frontal lobe

## Abstract

**Introduction:** Previous studies have shown that putative pheromones 4,16-androstadien-3-one (AND) and estra-1,3,5(10),16-tetraen-3-ol (EST) cause activation in the preoptic area/anterior hypothalamus in men and women. Sex differences in neural activation patterns have been demonstrated when participants are subject to pheromone stimulation; however, whether other compounds give rise to similar neural activity has not been completely investigated.

**Methods:** Twenty-nine young adults [16 female (21.3+/−0.54; mean yrs+/−SE), 13 male (22.85+/−0.42)] participated in a 3-block design, where participants were exposed to a scent (lavender), a synthetic male pheromone (4,16-androstadien-3b-ol; ALD), and a synthetic female pheromone (1,3,5(10),16-Estratetraen-3-ol; EST) via an automated olfactometer. Whole-brain, high-resolution (1.8mm^3^) functional MRI data from a Siemens Trio 3T MRI scanner were collected during all blocks. Five adults were excluded due to excessive movement. MANOVA analysis, a 2 × 3 multivariate model and analysis of 2×2 effects between sex and subsets of stimuli was done for activation over the whole brain and small volumes involved in olfaction.

**Results:** Exploratory analysis of 2×2 effects between sex and subsets of stimuli exhibited significant interactions when assessing activations over the whole brain, and small volumes involved in olfaction. The left and right frontal poles (LFP, RFP) shows significant interaction when assessing sex with lavender and EST for whole brain analysis. For small volume analysis, the right orbitofrontal cortex (ROFC) exhibited a sex with lavender and ALD interaction, and a sex with lavender and EST interaction was observed in the left inferior frontal gyrus (LIFG). Main effects of sex, stimulus, or interaction show no differences analyzed using a 2 × 3 multivariate model.

**Conclusion:** The study shows there is a sexually dimorphic response in the olfactory system to pheromones not previously studied. Scents like lavender do not have this same response. These distinct functional differences in activation patterns may be a result of neural development and maturation differences between sexes. Future studies should expand this pilot study and involve a younger demographic to accurately determine the age at which the olfactory response differentiates between males and females.

## Introduction

Many species can communicate through exogenous chemo-signals [1]. These chemo-signals can serve a variety of purposes including guiding sexual and social behavioral cues as well as survival aids in finding prey [2, 3]. Through extensive research, these chemo-signals were originally defined as pheromones – substances that were transmitted between two separate individuals of the same species that could influence behavioral or physiological changes in the receiving individual organism [4]. Since this time, researchers have attempted to shed light on the question of whether humans generate, and are affected by, pheromones.

Although it is still not fully clear whether humans produce and sense pheromones, there is evidence that chemosensory molecules elicit neural activity in humans who are exposed to them [5–7]. . For example, infants are attracted to chemo-signals that are secreted by the areola of the nursing mother [8]. Additionally, the structure of the human axilla facilitates the release of volatile compounds [9], and the flora and chemical content of the axilla are sexually dimorphic [10] suggesting that the axilla is likely a good site for the release of pheromone-like compounds.

One such pheromone-like compound, androstadienone (AND), has been found consistently in male axillary sweat [11, 12]. This compound, and estratetraene (EST), a molecule found in female human secretions, have been found to effect neural activity [13–15]. This has led investigators to propose AND, EST, and other similar steroid molecules as potential candidates for human pheromones [16, 17].

Whether humans actually produce putative pheromones has been a topic of debate [18]. Still, several studies have investigated and shown that exposure to AND affects the activity of the hypothalamus, orbitofrontal, piriform, fusiform, superior temporal gyri, and more recently, the amygdala [6, 19, 20]. Importantly, these areas are known to influence behavioral output and social and emotional processing [21–23], supporting the hypothesis that these molecules could have behavioral or physiological effects in receiving individuals.

Understanding how these signals are transmitted, received, and processed is further complicated by sexual dimorphism of the brain. Past research has shown significant differences in both volume and brain activity across both male and female individuals [24, 25]. For instance, chemo-signaling compounds, AND and EST, have been shown to cause similar activation patterns in individuals of the same sex [20].

While these studies investigated AND and EST, studies of animal models show that pheromones, or chemo signals, function with precise ratios of mixtures of compounds [26]. This suggests that chemo signals would naturally consist of more than AND or EST alone. Notably, more compounds than AND are found in the axillary sweat of humans, some of which are directly secreted by aprocrine sweat glands, others that are converted by bacterial flora [27]. These compounds likewise may act as chemo-signals that affect neural activity. Indeed, another similar molecule, Androstenol, was shown to elicit sexual-dimorphic neural activity in non-odor related areas of the brain [13]. Using functional magnetic resonance imaging (fMRI) protocols that measure brain activity in response to different inhaled compounds, the differences in chemo-signal induced brain activity between male and female participants can be shown.

Past studies have focused on AND or EST, thus the current study seeks to investigate the effects of Androsta-4,16-dien-3β-ol (Aloradine, ALD), a precursor of AND, as a potential human chemo-signal and to observe the differences in young adult brain activation when exposed to this chemo-signal molecule compared to EST.

## Methods

### Participants

Twenty-nine young, healthy adults (16 female [21.3 ± 0.54; mean years ± SE], 13 male [22.85 ± 0.42]) were recruited from the university and local communities. All participants were screened for and excluded if positive for any history of TBI, diagnosis of neurological condition, diagnosis of psychiatric condition, left-handed, non-native English speakers, or color blindness/deficient. Participants were also screened for MRI compatibility and were excluded if they had current permanent retainer/braces, pregnancy, prior injury to eye with metal object, any foreign metal implant in the body (e.g., pacemaker), or history of injury by metal object or foreign body. Patients were also screened for lavender allergies and excluded. The local Institutional Review Board approved all research protocols and participants gave written informed consent prior to participation.

### Study Design

A block design fMRI task was used, consisting of three blocks wherein participants were exposed to three different odorants: a lavender scent, a synthetic male pheromone (4,16-androstadien-3b-ol), and a synthetic female pheromone (1,3,5(10),16-Estratetraen-3-ol) via an automated olfactometer. The pheromones where diluted in propylene glycol to a concentration of 10mM, the mean detection threshold [5]. Before being placed in the scanner, participants were fitted with a nasal cannula with 5 m of tubing that extended through a wave guide in the wall between the MRI scanner room and the control room and connected to a custom olfactometer. Participants were instructed to breath normally through their nose. Odorants, contained in three separate airtight receptacles, were released to the participant by diverting the airflow through the specified receptacle on the olfactometer and through the cannula. Each block consisted of three alternating exposures of odorless air and one of the three odorants. The order of odorant presentation was pseudorandomized for each participant. To ensure jitter was achieved in the stimulus presentation, the segments of air preceding each odorant exposure were varied in duration at 30s, 35s, or 40s. Each exposure to the odorant lasted for 10s. Given the distance between the source of the odorant in the control room and the participant in the scanner, to ensure that participant was being exposed to the odorant during the acquisition, only the last five seconds of odorant exposure was used in the analysis. MRI data from exposure to odorless air was used as a baseline. To ensure participants were being exposed to just odorless air during baseline readings, and to maintain consistency with the duration of stimulus readings, the first five seconds in the 10 seconds preceding odorant release were used for baseline readings. Changes in odorant exposure were made without any visual or auditory cues to the participant.

### MRI Acquisition

All scans were acquired at the Brigham Young University MRI Research Facility on a Siemens Tim Trio 3T MRI scanner using a 32-channel receive-only head coil. All participants contributed both structural and functional images. Standard-resolution T1-weighted magnetization prepared rapid acquisition with gradient echo (MPRAGE) structural images were acquired with the following parameters: TR = 20 ms, TE = 4.92 ms, flip angle = 25°, FoV = 256 × 256, voxel size = isotropic 1 mm, slices = 192 interleaved. High-resolution multi-band echo-planar T2*-weighted functional images were acquired using a pulse sequence with the following parameters: multi-band factor = 8, TR = 875 ms, TE = 43.6 ms, flip angle = 55°, acquisition matrix = 100 ×100 mm, voxel size = isotropic 1.8 mm, slices = 72 interleaved.

### Pre-processing

Analyses were conducted by a blinded researcher with the software packages dcm2niix [28], Analysis of Functional NeuroImages [29], convert 3D [30], and visualized in Mango [31].

T1- and T2*-weighted DICOMs were first converted to NIfTI files, and then the T1-weighted file was aligned to the EPI volume containing the smallest number of outlier voxels using a rigid transformation with a Local Pearson Correlation cost function. The rigid-alignment calculations were concatenated with both those of a nonlinear diffeomorphic transformation, between the T1 image and a template in MNI space, and the T2* volume registration calculations in order to move all T2* data into template space using a single interpolation. The T2* data were then scaled by the mean signal at each voxel. Single-subject regressions were modelled with a generalized-least-squares fit that utilized a nonlinear residual-maximum-likelihood estimate to best fit the autoregressive-moving-average model. Included in the model were a nuisance regressor utilizing the white matter time series, centered motion regressors for six degrees of freedom, and regressors which corresponded to the last five seconds of each odorant trial that were modeled as a block function convolved with the canonical hemodynamic response function.

Baseline for the model consisted of time periods when participants were exposed to odorless air. Any volumes with excessive movement (>0.3° translation) and with a large percentage of outlier voxels (>10%) were excluded, as was the preceding volume. Exclusion criteria for the group-level analysis was determined *a priori* to exclude any participant with more than 10% of their total volumes being censored due to outlier signal or motion of participant during the scan (the censor file was the result of multiply a binary matrix which indicated which volumes contained outlier signal in >10% of their voxels by a binary matrix indicating which volumes had significant movement events); this resulted in three women and two men being excluded from the group-level analysis due to excessive movement. Thus, a total of 24 subjects were included in final analysis.

For group-level analyses, an inclusion mask was constructed by multiplying a gray matter mask with an intersection mask; briefly, the intersection mask is the result of mapping where each participant had sufficient T2*-weighted signal for analyses. This is done to account for signal fallout. Following updated recommendations [32, 33] autocorrelation function parameter estimates were then derived from the blurred (full-weight, half-maximum Gaussian, blur size = 2 × voxel dimensions) model residuals, and these parameter estimates were used in Monte Carlo noise simulations to determine thresholding criteria. Thresholding criteria were calculated for both whole-brain and small-volume analyses inclusion masks, where the small-volume consisted of the bilateral amygdalae, piriform/entorhinal, and orbitofrontal regions. The threshold criteria used were nearest neighbors = 1, p < .001, α = .05, and cluster size = 19 (whole-brain) and 8 (small-volume) voxels. For any clusters surviving multiple-comparison thresholding, mean β-coefficients were then extracted for subsequent analyses. All scripts used in these pre-processing steps as well as for the group-level analyses are available at https://github.com/nmuncy/Pheromone.

We performed a MANOVA analysis to test the between-subjects effect for each of the areas of activation. In order to determine if men and women differentially processed sex-specific pheromones, a 2 × 3 multivariate model was used to potentially identify regions which demonstrated differential signal, where sex was considered a between-subject factor and stimulus type (Lavender, ALD, and EST) a within-subject factor. Analysis of 2×2 effects between sex and subsets of stimuli was done for activation over the whole brain and small volumes involved in olfaction.

## Results

### Whole-brain analyses

In the 2×3 multivariate model, no region had detectable differences for a main effect of sex, a main effect of stimulus, or an interaction thereof. An exploratory 2×2 contrast of sex with lavender and EST did, however, show activation differences by sex in the left (*F*(2,21) = 18.040, *p* < 0.001;Wilk’s Λ = 0.368, *η*^2^ = 0.632;*observed power* = 0.999;) and right (*F* (2,21) = 7.008, *p* < 0.005;Wilk’s Λ = 0.600, *η*^2^ = 0.4003; *observed power* = 0.886;) frontal pole (**Error! Reference source not found.**, A-B; **Error! Reference source not found.**). Follow up univariate analysis of between-subject factors indicate a treading difference in activation response to EST between sexes for the LFP (*F*(1,22) = 3.281, *p* = 0.084; *η*^2^ = 0.130;*observed power* = 0.41) but not RFP *F*(1,22) = 2.086, *p* = 0.163;*η*^2^ = 0.087;*observed power* = 0.282.

**Figure 1:**
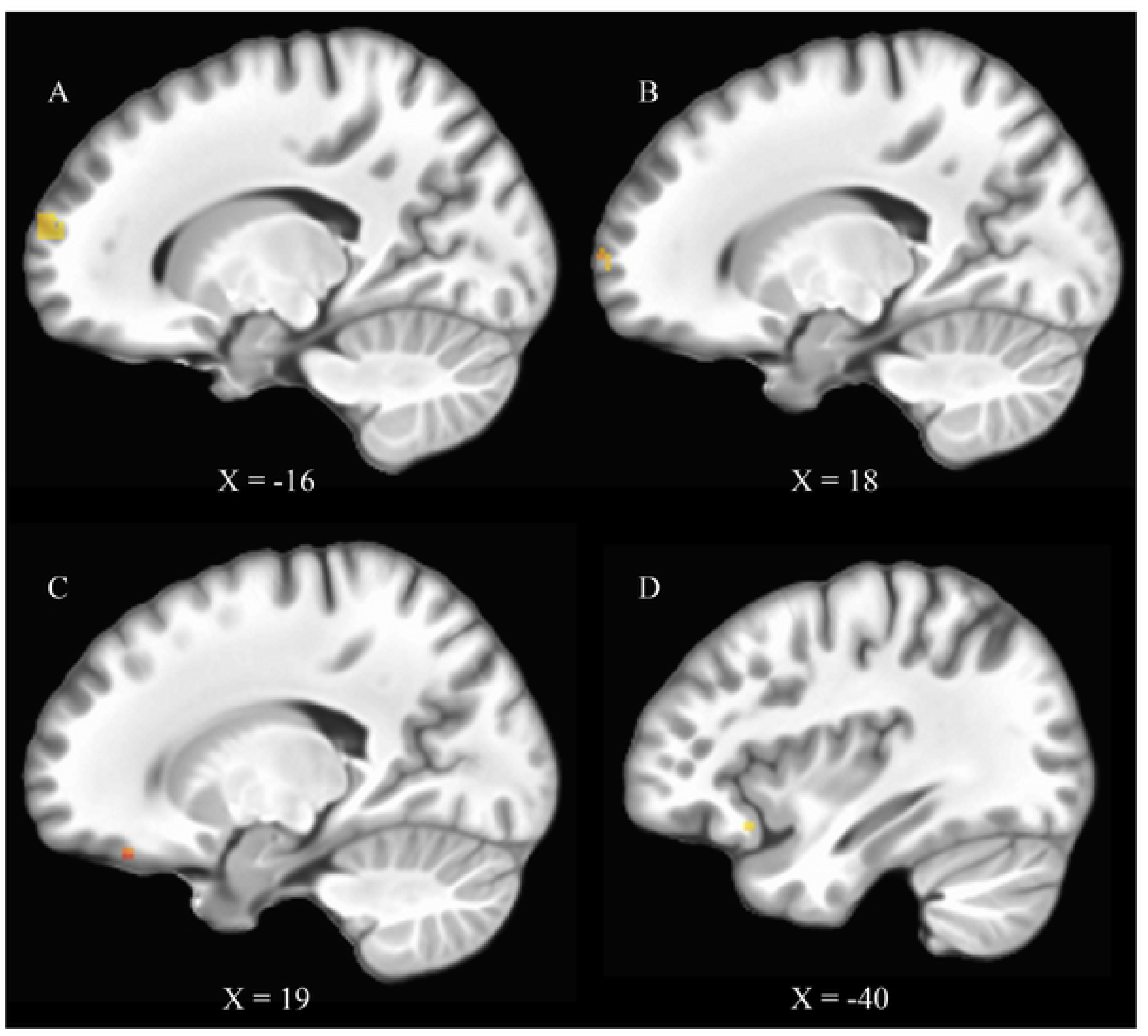
Top, clusters surviving whole-brain thresholding in left (A) and right (B) frontal pole regions for a sex × Lavender, EST interaction. Bottom, clusters surviving a small-volume correction. C the right orbitofrontal cortex showed a sex × Lavender, ALD interaction, and D the left inferior frontal gyrus showed a sex × Lavender, EST interaction.

### Small-volume analyses

As smaller regions are involved in olfaction, such as the piriform cortex, and these same regions may be smaller than the multiple-comparison correction, a small-volume analysis was conducted on the bilateral amygdalae, piriform, and orbitofrontal cortices using the same 2 × 3 multivariate model as above. As before, no cluster was detected which showed a main effect of sex, stimulus, or interaction thereof, but exploratory analyses revealed two additional clusters: the right orbitofrontal cortex showed a sex × Lav, ALD interaction (*F*(2,21) = 7.309, *p* < 0.004; *Wilk’s Λ* = 0.590, *η*2 = 0.4104; *observed power* = 0.899) and the left inferior frontal gyrus showed a sex × Lav, EST interaction (*F*(2,21) = 16.889, *p* < 0.001; *Wilk’s Λ* = 3.83, *η*2 = 0.6166; *observed power* = 0.999) (**Error! Reference source not found.**, C-D, respectively, **Error! Reference source not found.**). Follow-up univariate analysis indicated a significant difference in activity of the LIFG between sexes exposed to EST (*F*(1,22) = 32.021,*p* < 0.0001;*η*^2^ = 0.593;*observed power* = 1.00) but only a trending toward significance for activity of the ROFC between sexes exposed to ALD (*F*(1,22) = 3.328, *p* = 0.082;*η*^2^ = 0.131;*observed power* = 0.415). Analysis of the estimated marginal means revealed a significant difference between sexes within the LIFG when exposed to EST (Figure 2E, **Error! Reference source not found.**H).

**Figure 2:**
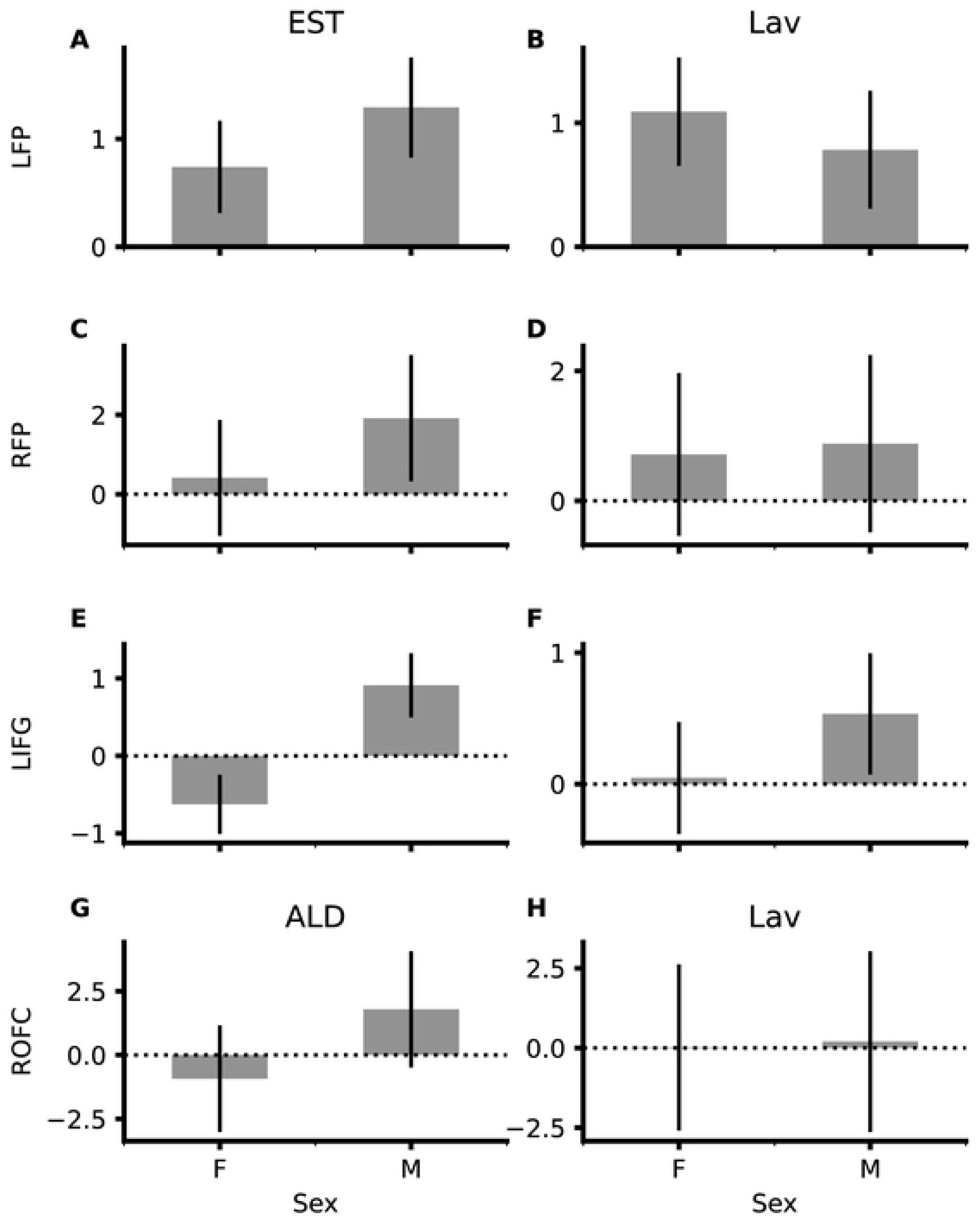
Estimated Marginal Means by subjects’ sex in the left frontal pole (A, B), right frontal pole (C, D), left inferior frontal gyrus (E, F), and the right orbitofrontal cortex (G, H), when subjects where exposed to EST (A,C,E), ALD (G) or lavender (B,D,F,H). *Error reported as CI-95.*

## Discussion

Our study shows that a sexually dimorphic response to chemo-sensory signals exists in the olfactory system but not to normal scents such as lavender. This confirms similar findings in previous reports, that androgen- and estrogen-related chemo-signals differentially affect males and females [19]. It’s important to note however, that previous studies have investigated only a narrow selection of possible chemo-sensory stimulants, restricting their selection to AND and andostradiol. Our newly selected compounds were chosen because of associations, which previous studies made, with the effects of the compounds in animals [34]. Here, we utilized 4,16-androstadiene-3-ol (ALD), an additional type of androgen and were able to elicit a sexual dimorphic neural activation profile similarly observed in previous studies. This suggests that these chemosensory-induced neural activation patterns are not dependent on one or two specific compounds but may exhibit similar patterns of activity for similar chemical compounds within same-gendered participants. Future studies should use various moieties of AND and EST to elucidate the necessary components of these steroids that induce similar neural activation patterns to better understand how a range of steroids can affect the human brain.

Currently, the OFC and the PFC are two ROIs that show consistent activation across studies [15, 19]. These ROIs give rise to social and emotional processing, and together, these may be associated with a sense of attractiveness [35]. Studies have hypothesized links between activation of these brain areas and social and emotional processing [6, 21], on the idea that, like pheromones, these chemosensory signals may function as behavioral cues between individuals within the same species. Since the development of emotional and social processing is multifactorial, and not simply a result of genetics, these distinct functional differences in activation patterns could be a consequence of neural development and maturation, such that observed activation patterns within same-gendered participants are age dependent. Future studies should involve a younger demographic to accurately determine the age at which the observed olfactory response differentiates between young and adult males and females.

Aside from OFC and PFC, our results highlight activation differences in areas associated with a set of cognitive processes. The bilateral medial prefrontal cortex can be activated during memory and decision making [36], the inferior frontal pole is shown to activate in activities associated with forward thinking and planning [37], and the right superior frontal sulcus plays a role in impulse control [38]. Furthermore, this includes emotional processing for the PFC and OFC [23], sensory for the MFS, memory for the lSFS [39], attention control in the rMFS [40], and motor control processing for the raSFS. Taken together, it would appear that the neural activation of these compounds are associated with complex neural processing. Future work of interest could observe the activational differences in participants during tasks that recruit these areas while exposed to chemosensory compounds.

Our data showed a differentiation of activation patterns in the following areas: LFP (sex × Lavender, EST interaction), RFP (sex × Lavender, EST interaction), ROFC (sex × Lavender, ALD interaction), and LIFG (sex × Lavender, EST interaction). The frontal poles are considered part of the pre-frontal cortex; thus, activation of the LFP, RFP, and ROFC is consistent with data in prior studies [15, 19]. LIFG has been reported in the literature as playing a role in implementing reappraisal strategies [41] as part of social and emotional processing, as well as having a unique connectivity to piriform cortex of olfactory processing pathway, [42] thus it is thought to be part of secondary olfactory areas [43].

One outcome of particular interest in this study is that chemosignal-induced activation was located primarily cortical, including the orbitofrontal pathway with its connection to the amygdala and involvement in emotional relay [44], in addition to the left superior frontal sulcus, associated with working memory [39]. This pattern of activation suggests that simply smelling the compounds may induce activation in emotional or memory centers. This needs to be studied in greater detail, as our experiments did not directly target these cortical functions. Follow-up studies of the effects of the exposure of these compounds on participants’ memory may cultivate new information in this sphere. However, moving forward, our results importantly diversify and expand new compounds can be used to further elucidate the influence of inducing neural activity in the areas observed in the context of emotional relationships, mental disease, and possibly sexual trauma.

Our study does have some limitations. It is important to note the 10mM concentration is considered a “high” concentration, as taken from [5], and that this concentration is above physiological relevance, but was shown to give rise to the dimorphic effects seen in other studies. A previous study showed differences in results based on compound concentration [45], however, future studies need to explore the effects of varying concentrations at more physiologically-relevant concentrations. Another important consideration is that during the testing period, participants were prompted to breathe normally. As people breathe differently (with different respiratory rates and volumes) this could potentially affect the amount of compound that each participant was exposed to even with uniform concentration. Lastly, the number of subjects used in the study is low, 24. It’s possible that given the variability in the data collected, increased subject variance may add to the noise of our findings, decreasing the generalizability of our findings. Despite this, the observed power in our analysis suggests our results effectively found differences where they existed and lays groundwork as a pilot for future studies with increased subject numbers.

Further, previous studies have shown not only that responses to chemosensory compounds exhibit sexual dimorphism, but there are also differences in responses based on sexual-orientation. Since this was a pilot study with the focus of obtaining a baseline response to the pheromones, we utilized an opportunity sample based on previously approved recruiting measures, which did not include screening for sexual-orientation. This leaves room for future investigation, as prior studies have shown that homosexual men and women tended to respond opposite to their heterosexual counterparts. While we still observed activation differences under the current assumption of all heterosexual participants, it will be important to consider the factor in continuing studies.

Our study demonstrates that other specific chemosensory compounds exist in addition to AND and EST as previously shown. All capable of evoking sexually-dimorphic and neuroanatomically specific activation of various brain regions across participants. This study provides motivation for further investigation into chemosensory-evoked patterns neural activation by varying chemical compounds, at varying concentrations, and under task-oriented paradigms. Elucidating the answer to these topics will help deepen our understanding of the role that chemosensory compounds play in humans.

**Table 1:** 2×2 Multivariate Analysis between sex and selected chemo-signals. Coordinates and statistics for whole-brain and small-volume analyses. Size = cluster size, X, Y, Z = center of mass.

| Analysis | Contrast | Cluster | F-stat | Sig | Effect | Observed<br>Power | Size | X | Y | Z |
| --- | --- | --- | --- | --- | --- | --- | --- | --- | --- | --- |
| Whole brain | Sex × Lav, EST | L. Frontal Pole | 18.040 | 0.000 | 0.6321 | 0.999 | 45 | 16 | 67 | 17 |
|  |  | R. Frontal Pole | 7.008 | 0.005 | 0.4003 | 0.866 | 19 | 18 | 69 | 6 |
| Small volume | Sex × Lav, ALD | R. Orbitofrontal<br>Cortex | 7.309 | 0.004 | 0.4104 | 0.899 | 8 | 19 | 39 | -22 |
|  | Sex × Lav, EST | L. Inferior Frontal<br>Gyrus | 16.889 | 0.000 | 0.6166 | 0.999 | 8 | -36 | 23 | -18 |

**Table 2:**
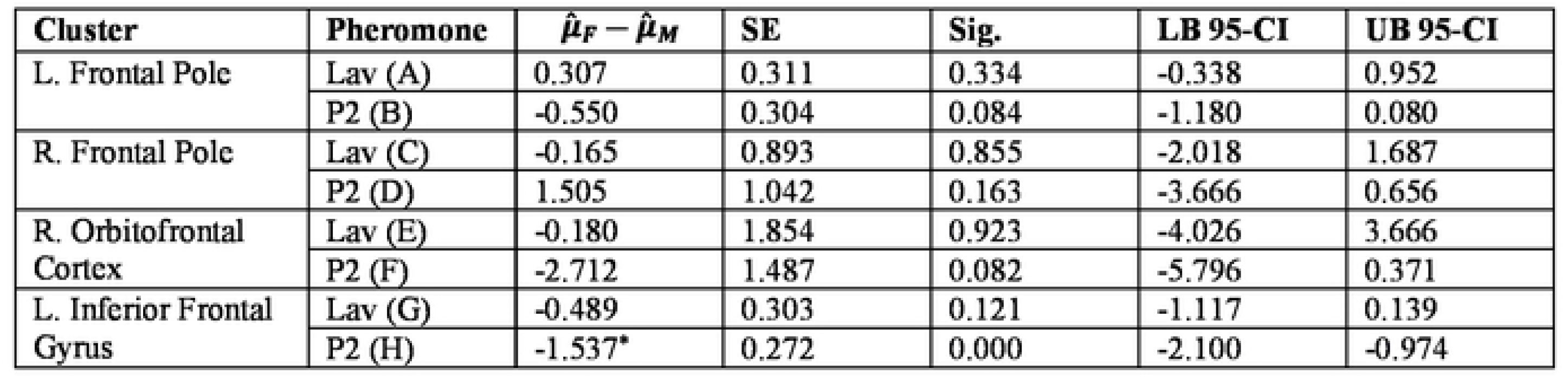
Pairwise comparisons per cluster.

## Acknowledgments

The authors would like to thank the following entities for their generous support of this research: Brigham Young University, College of Life Sciences, Mentoring Environment Grant; Brigham Young University, School of Family Life, Gerontology Program; Brigham Young University, Magnetic Resonance Imaging Research Facility Seed Grant.

